# Integer linear programming outperforms simulated annealing for solving conservation planning problems

**DOI:** 10.1101/847632

**Authors:** Richard Schuster, Jeffrey O. Hanson, Matt Strimas-Mackey, Joseph R. Bennett

**Author notes:** Corresponding author: Department of Biology, 1125 Colonel By Drive, Carleton University, Ottawa ON, K1S 5B6 Canada.

## Abstract

The resources available for conserving biodiversity are limited, and so protected areas need to be established in places that will achieve objectives for minimal cost. Two of the main algorithms for solving systematic conservation planning problems are Simulated Annealing (SA) and Integer linear programming (ILP). Using a case study in British Columbia, Canada, we compare the cost-effectiveness and processing times of SA versus ILP using both commercial and open-source algorithms. Plans for expanding protected area systems based on ILP algorithms were 12 to 30% cheaper than plans using SA. The best ILP solver we examined was on average 1071 times faster than the SA algorithm tested. The performance advantages of ILP solvers were also observed when we aimed for spatially compact solutions by including a boundary penalty. One practical advantage of using ILP over SA is that the analysis does not require calibration, saving even more time. Given the performance of ILP solvers, they can be used to generate conservation plans in real-time during stakeholder meetings and can facilitate rapid sensitivity analysis, and contribute to a more transparent, inclusive, and defensible decision-making process.

## Introduction

Area-based systematic conservation planning aims to provide a rigorous, repeatable, and structured approach for designing new protected areas that efficiently meet conservation objectives (Margules and Pressey 2000). Historically, spatial conservation decision-making often evaluated parcels opportunistically as they became available for purchase, donation, or under threat (Pressey et al. 1993, Pressey and Bottrill 2008). Although purchasing such areas may improve the status quo, such decisions may not substantially and cost-effectively enhance the long-term persistence of species or communities (Joppa and Pfaff 2009, Venter et al. 2014). Systematic conservation planning, on the other hand, involves framing conservation planning problems as optimization problems with clearly defined objectives (e.g. minimize acquisition cost) and constraints. These optimization problems are then solved to obtain candidate reserve designs (termed solutions), which are used to guide protected area acquisitions and land policy (Schwartz et al. 2018). Due to the systematic, evidence-based nature of these tools, they can help contribute to a transparent, inclusive, and more defensible decision-making process (Margules and Pressey 2000).

Today, Marxan is the most widely used systematic conservation planning software, having been used in 184 countries to design marine and terrestrial reserve systems (Ball et al. 2009). Although Marxan supports several algorithms for solving conservation planning problems, most conservation planning exercises use its implementation of simulated annealing (SA), an iterative, stochastic metaheuristic algorithm for approximating global optima of complex functions with many local optima (Kirkpatrick et al. 1983). By conducting thousands of individual runs, each with millions of iterations, Marxan aims to generate solutions that are near-optimal. One of the reasons why Marxan uses SA instead of integer linear programming (ILP), is that ILP was not well suited to solve problems with nonlinear constraints and penalties, such as problems trying to create spatially compact or connected solutions (i.e. compactness and connectivity goals). However, the SA approach provides no guarantee on solution quality. As a consequence, conservation scientists and practitioners have no way of knowing if their solutions are highly suboptimal.

In a recent simulation study, Beyer et al. (2016) found that Marxan with simulated annealing can deliver solutions that are orders of magnitude below optimality. They compared Marxan to integer linear programming (ILP) (Dantzig 2016), which minimizes or maximizes an objective function (a mathematical equation describing the relationship between actions and outcomes) subject to a set of constraints and conditional on the decision variables (the variables corresponding to the selection of actions to implement) being integers (Beyer et al. 2016). Unlike metaheuristic methods such as SA, prioritization using ILP will find the exact optimal solution or can be instructed to return solutions within a defined distance from optimality. Some have argued that ILP algorithms are well-suited for solving conservation planning problems (Cocks and Baird 1989, Underhill 1994, Rodrigues and Gaston 2002), but until recent advances in computational capacity and algorithms, it has been impossible to solve the Marxan-like systematic conservation planning problems with ILP for large problems (Beyer et al. 2016).

Here we compare integer linear programming with simulated annealing (i.e. Marxan) for solving systematic conservation planning problems using real-world data from Western North America. We found that ILP generated high quality solutions 1,000 times faster than simulated annealing that could save over $100 million (or 13%) for realistic conservation scenarios when compared to solutions obtained from simulated annealing. These results also hold true for problems aiming for spatially compact solutions. Our findings open up new possibilities for scenario generation to quickly explore and compare different conservation prioritization scenarios in real-time.

## Material and Methods

### Study area

We focused on a 27,250 km^2^ portion of the Georgia Basin, Puget Trough and Willamette Valley of the Pacific Northwest region spanning the US and Canada, corresponding to the climate envelope indicative of the Coastal Douglas-fir (CDF) Biogeoclimatic zone in southwestern British Columbia (Meidinger and Pojar 1991) (Appendix S1: Figure S1). Land cover in the region is diverse, with approximately 57% of the land in forest, 8% as savanna or grassland, 5% in cropland, 10% being urban or built and the rest in wetland, water or barren.

### Biodiversity data

We used species distribution models for 72 bird species as our conservation features at a 1-ha grid cell resoltuion (Supplementary Table 1). The distribution models were based on data from eBird, a citizen-science effort that has produced the largest and most rapidly growing biodiversity database in the world (Hochachka et al. 2012, Sullivan et al. 2014). From the 2013 eBird Reference Dataset (http://ebird.org/ebird/data/download) we used a total of 12,081 checklists in our study area, then filtered these checklists to retain only those from March – June to capture the breeding season, <1.5 hours in duration, <5 km travelled, and a maximum of 10 visits to a given location to improve model fit. Sampling locations <100 m apart were collapsed to one location, yielding 5,470 checklists from 2,160 locations, visited from 1-10 times and 2.53 times on average. The R package unmarked (version 0.9-9; Fiske and Chandler 2011) provided the framework for all species distribution models, which necessarily include two parts: occupancy and detection (Mackenzie et al. 2002). For further details on biodiversity data see (Rodewald et al. 2019).

**Table 1.**
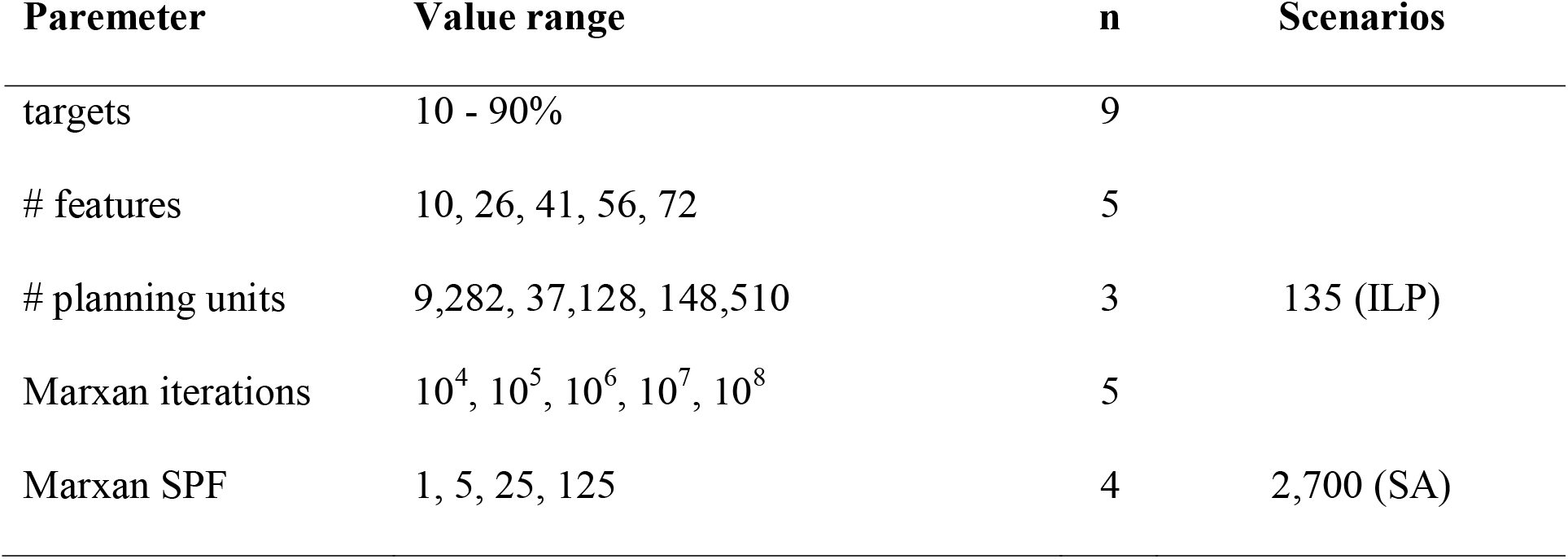
Scenarios investigated in our analysis. The total number of scenarios tested for both Gurobi and SYMPHONY are 135. For Marxan analysis, we included calibration steps as well, which brought the total number of scenarios to 2700 for that algorithm.

### Cadastral layer and land cost

We incorporated spatial heterogeneity in land cost (Ando et al. 1998, Polasky et al. 2001, Ferraro 2003, Naidoo et al. 2006) in our plans by using cadastral data and 2012 land value assessments from the Integrated Cadastral Information Society of BC. This process resulted in 193,623 polygons for BC which were subsequently used as planning units (Schuster et al. 2014). Cadastral data, including tax assessment land values from Washington State came from the University of Washington’s Washington State Parcel Database (https://depts.washington.edu/wagis/projects/parcels/; Version: StatewideParcels_v2012n_e9.2_r1.3; Date accessed: 2015/04/30), as well as San Juan County Parcel Data with separate signed user agreement. The combined cadastral layer included 1.92 million polygons. Cadastral data, including tax assessment land values from Oregon State had to be sourced from individual counties, which included Benton, Clackamas, Columbia, Douglas, Lane, Linn, Marion, Multnomah, Polk, Washington and Yamhill. The combined cadastral layer for Oregon included 605,425 polygons. We converted the polygon cost values to 1-ha raster cells for consistency with the biodiversity data by calculating area weighted mean values of cost per raster cell.

### Spatial prioritization

We compared ILP and SA for solving the minimum set spatial prioritization problem (Ball et al. 2009). In this formulation, the landscape is divided into a set of discrete planning units. Each planning unit is assigned a socioeconomic cost (here we use the assessed land value) and a conservation value for a set of features that we wish to protect (here the occupancy probability for a set of species). Finally, we define representation targets for each species as the amount of habitat we hope to protect for that species. The goal of this prioritization problem is to optimize the trade-off between conservation benefit and socioeconomic cost (McIntosh et al. 2017). Achieving this goal involves finding the set of planning units that meets the conservation targets for the minimum possible cost (i.e. min cost: such that conservation value ≥ target). Details on the Marxan problem formulation can be found in Ball et al. (2009) and the ILP formulation in Beyer et al. (2016). Three key parameters that are important for Marxan analysis, which we also use here are: species penalty factor, number of iterations, and number of restarts (Ardron et al. 2010). Briefly, the species penalty factor is the penalty given to a reserve system for not adequately representing a feature, the number of iterations determines how long the annealing algorithms will run, and the number of restarts determines how many different solutions Marxan will generate. For all scenarios, we used 1 km^2^ planning units, generated by aggregating the species and cost data to this coarser resolution from the original 1-ha cells. Aggregation was accomplished by taking the sum of cost data and the mean of species data for all 1-ha cells within the larger 1 km^2^ cells.

### ILP solvers (commercial vs open source)

A variety of ILP solvers currently exist, and both commercial and open source solvers are available. All solvers yield optimal solutions to ILP problems, but there are substantial differences in performance (i.e. time taken to solve a problem) and in the size of problems that can be solved (Lin et al. 2017). For the purposes of performance testing we opted for one of the best commercial solvers currently available, Gurobi (Gurobi Optimization Inc. 2017). In a recent benchmark study, Gurobi outperformed other solver packages for more complex formulations and a practical use-case (Luppold et al. 2018). To investigate solver performance of packages that are freely available to everyone, we also tested the open source solver SYMPHONY (Ralphs et al. 2019). Both Gurobi and SYMPHONY can be used from R. For Gurobi we used the R package provided with the software (Gurobi version 8.1-0) and for SYMPHONY the Rsymphony package (version 0.1-28; Harter et al. 2017). We used the prioritizr R package to solve ILP problems for both Gurobi and SYMPHONY solvers (Hanson et al. 2019).

### Scenarios investigated

We investigated a range of scenarios that were computationally feasible for this study. For both Marxan and prioritzr we created the following range of scenarios: i) vary conservation targets between 10 and 90% protection of features in 10% increments (9 variations), using ii) 10 – 72 species/features (5 variations) as targets, and iii) with spatial extents of 9,282, 37,128, and 148,510 planning units (3 variations), resulting in a total of 135 scenarios created (Table 1). For Marxan, we also varied two additional parameters, i) the number of iterations ranged from 10^4^ to 10^8^ (5 variations) and ii) species penalty factors (SPF) of 1, 5, 25, and 125 were explored (4 variations, roughly spanning two orders of magnitude) for a total of 2,700 scenarios investigated in Marxan (Table 1). As the processing time for the most complex problem in Marxan (90% target, 72 features, 148,510 planning units, 10^8^ iterations) was >8 hours, we restricted the full range of scenarios to those mentioned above. The maximum number of planning units we used is within the range of previous studies using Marxan (e.g. Venter et al. 2014; Runge et al. 2016), although using more than 50,000 planning units with SA is discouraged without extensive parameter calibration, as near optimal solutions will be hard to find for problems of that size (Ardron et al. 2010).

As systematic conservation planners often aim for spatially compact solutions to their problems, we also investigated a range of scenarios using a term called boundary length modified (BLM), which is used to improve the clustering and compactness of a solution (McDonnell et al. 2002). We randomly selected a 225 × 225 pixel region of the study area to generate a problem with 50, 625 planning units, the maximum recommended for Marxan. After initial calibration we set the number of features/species to 72, SPF to 25 and number of iterations for Marxan to 10^8^. We varied targets between 10 and 90% protection of features in 10% increments, and used the following BLM values: 0.1; 1; 10; 100; 1,000 for a total of 45 scenarios.

All analyses were conducted on a desktop computer with an Intel Core i7-7820X Processor and 128 GB RAM running Ubuntu 18.04 and R v 3.5.3. All data, scripts and full results are available here: https://osf.io/my8pc/

## Results

ILP algorithms (Gurobi, SYMPHONY) outperformed SA (Marxan) in terms of their ability to find minimal cost solutions across all scenarios that met conservation targets. Through finding optimal solutions, using ILP resulted in cost savings ranging from 0.8% to 4,369% (median 72.7%). When we restricted results to only take into account calibrated Marxan scenarios (number of iterations > 100,000 and species penalty factor 5 or 25), the range of savings was reduced to 0.8% to 52.5% (median 12.6%, Appendix S1: Figure S2). For example, at the 30% protection target ILP solvers resulted in solutions that were $144 million cheaper than SA (Figure 1a). With these savings an additional 3,039 ha could be protected (53,934 ha vs 50,895 ha) using an ILP algorithm by raising the representation targets until the cost of the resulting solution matched that of the Marxan solution using SA. In general, SA performed reasonably well at smaller problem sizes, fewer planning units and features and low targets, but as the problem size and complexity increased SA was less consistent in finding good solutions (Appendix S1: Figure S2). Cost profiles across targets, number of features and number of planning units are shown in Appendix S1: Figures S3-5.

**Figure 1.**
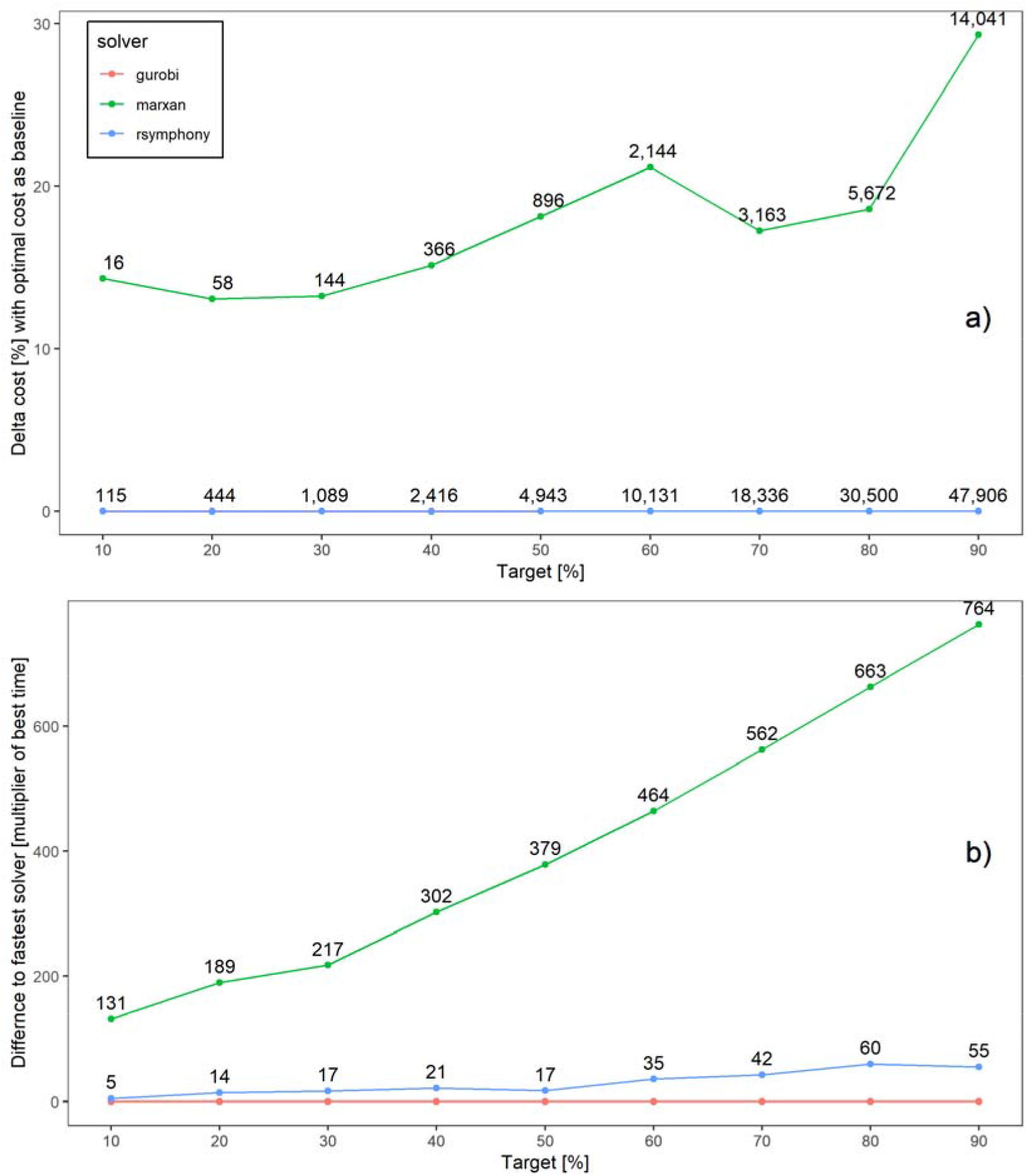
Solution cost and time comparisons. a) The lines represent costs compared to the Gurobi cost baseline. The numbers on the blue line represent total cost of a solution in million $ and the numbers on the green line represent how much more expensive, again in million $, the SA/Marxan solution is compared to the ILP solutions. b) Time to solution comparisons between solvers. Marxan parameters used are: 72 features, 148,510 planning units, 10^8^ iterations, using mean cost and time. Note that in a) gurobi (red) and Rsymphony (blue) yielded optimal solutions for all target values and so their lines are plotted exactly on top of each other.

The shortest processing times were achieved using the prioritizr package and the commercial solver Gurobi, followed by prioritizr and the open source solver SYMPHONY, and lastly Marxan (Figure 1b). Gurobi had the shortest processing times across all scenarios investigated, SYMPHONY tied with Gurobi in some scenarios and took up to 78 times longer than Gurobi in other scenarios (mean = 14 times, Appendix S1: Figure S6), and Marxan took between 1.8 and 1995 times longer than Gurobi (mean = 281 times, Appendix S1: Figure S7). The longest processing times for Gurobi, SYMPHONY and Marxan for a single scenario were 40 seconds, 31 minutes, and 8 hours respectively. For the most complex problem (i.e. targets = 90%, 72 features; 148,510 planning units), Marxan calibration across the 5 number of iterations and 4 species penalty factor values took a total of 5 days 7 hours, compared to 30 seconds using Gurobi and 28 minutes using SYMPHONY. Time profiles across targets, number of features and number of planning units are shown in Appendix S1: Figures S8-10.

ILP algorithms (Gurobi, SYMPHONY) also outperformed SA (Marxan) when using a BLM to achieve compacter solutions. This was true for objective function values (Figure 2a) as well as for processing times (Figure 2b). Through finding optimal solutions, using ILP resulted in objective function values 5.65 to 149% (mean 22.7%) lower than SA values. Gurobi was the fastest solver to find solutions to problems including BLM in 44 of 45 scenarios, in one case SYMPHONY was faster. SYMPHONY outperformed Marxan in 44 of 45 scenarios, and took on average 13.7 times as long as Gurobi to find a solution (range −0.31 to 42.6). Marxan was never faster than Gurobi and took on average 104.6 times as long as Gurobi to find a solution (range 3.09 to 190.8). An example of the spatial representation of the solutions for a 10% target is shown in Appendix S1: Figure S11.

**Figure 2.**
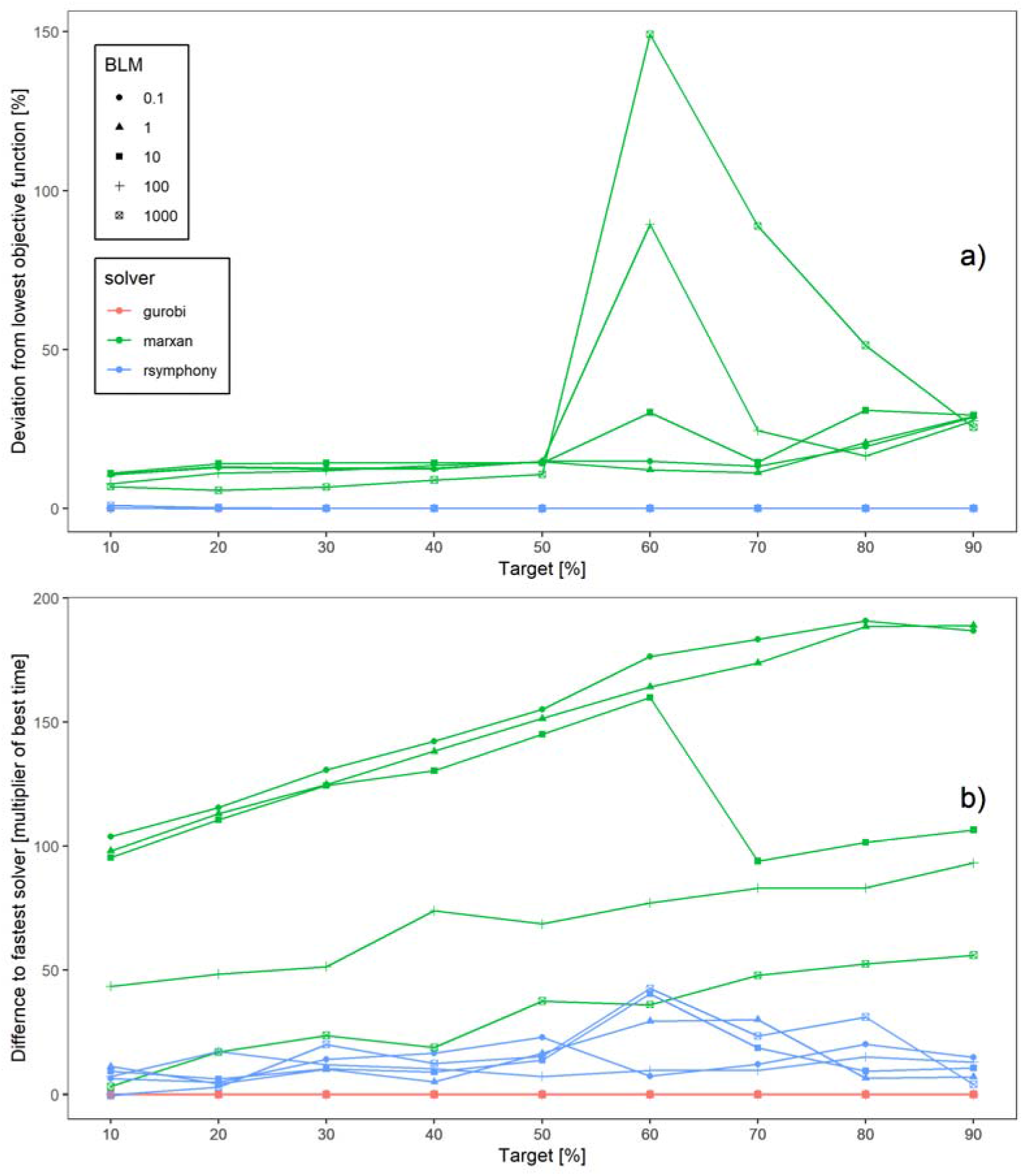
Objective function value and time comparisons using a boundary penalty to achieve spatially compact solutions. a) Deviation from lowest objective function value for solvers used and over a range of boundary penalty or boundary length modifier values (BLM); zero deviation indicates optimal solution. b) Time to solution comparisons between solvers and across BLM values. Note that in a) gurobi (red) and Rsymphony (blue) yielded optimal solutions for all target values and so their lines are plotted exactly on top of each other.

## Discussion

We found that ILP algorithms outperformed SA both in terms of cost-effectiveness and processing times, even when including non-linear problem formulations, when planning for spatially compact solutions. There have been calls for using ILP in solving conservation planning problems in the past (Underhill 1994, Rodrigues and Gaston 2002), but we are now at a point where making this switch is both advisable and computationally feasible. Our study provides a systematic test case, using real world data to build on the findings of (Beyer et al. 2016) and show that their results hold for a realistic case study. We further expanded the scope of testing to include assessed land values in order to give estimates of how much better optimal solution can perform in terms of cost savings, compared to SA solutions. Finally, we showcase that even open source ILP solvers are much faster than SA algorithms as implemented in Marxan, which is very encouraging for non-academic user that would otherwise have to buy Gurobi licenses (Gurobi is free for academic use). The combination of the superior performance findings by both (Beyer et al. 2016) and this study indicates that ILP approaches should be strongly considered as improvements for minimum set conservation planning problems, currently solved using SA.

One practical advantage of using ILP over SA is that the analysis does not require parameter calibration. Unlike ILP, parameter calibration is a crucial task in every Marxan/SA project and the species penalty factors, number of SA iterations, and number of SA restarts must be calibrated improve solution quality (Ardron et al. 2010). This task can be very time consuming, especially for larger problems (e.g. 50,000 planning units). Ideally all possible combinations of parameters should be explored, but this further increases processing time. For instance, exploring three different parameter values would result in 27 different scenarios to explore (i.e. 3 × 3 × 3). Although we omitted calibration runs prior to finalizing and presenting results in this study, the parameter calibration step took several days for the most complex problem we investigated in this study. Yet none of this calibration time is necessary using ILP. An added benefit is that the somewhat subjective process of setting values for these three parameters can be eliminated using ILP as well.

Recommended practices for Marxan analyses caution against using SA for conservation planning exercises with more than 50,000 planning units (Ardron et al. 2010). Such large-sized problems have occurred in the past and, as increasingly high resolution data become available, may become more common in the future (e.g. Venter et al. 2014; Runge et al. 2016). Unlike SA, ILP/prioritizr can solve problem sizes with more than one million planning units (Hanson 2018, Schuster et al. 2019). Realistically, as problem sizes grow beyond what was intended for Marxan/SA projects, ILP will run into problems solving very large problems (>1 million planning units) that include non-linear constraints, such as optimizing compactness or connectivity, as those problem formulations need to be linearized for ILP to work. There is the potential to use nonlinear integer programming for more complex problems in the future though (Grossmann 2002, Lee and Leyffer 2011). Whether ILP would also outperform SA for more complex problem formulations, such as dynamic problems or problems with multiple objectives, still needs to be explored. Potential solutions would be to linearize the problem, or incorporate algorithms like Mixed Integer Quadratically Constrained Programming (Franco et al. 2014).

Finally, we argue that another strength of ILP solvers, especially Gurobi, is that they can be used to quickly explore and compare different conservation prioritization scenarios in real-time. This ability could be used to great advantage during stakeholder meetings, to explore various scenarios and undertake rapid sensitivity analysis.

## Conclusion

ILP algorithms substantially outperform SA as used in minimum set systematic conservation planning, both in terms of solution cost, as well as in terms of time required to find near optimal or optimal solutions. Using an ILP algorithm, as implemented in the R package prioritizr, has the added benefit that users do not need to worry about or set parameters such as species penalty factors or number of iterations, which significantly reduces the time a user spends on finding suitable values for these parameters. Given the potential ILP is showing for conservation planning, we recommend users consider adding this modified approach to solving systematic conservation planning problems.

## Acknowledgements

RS is supported by a Liber Ero Fellowship and Environment and Climate Change Canada (ECCC), JOH by ECCC, MSM by endowments at the Cornell Lab of Ornithology, and JRB by Natural Sciences and Engineering Research Council of Canada and ECCC. We thank W. Hochachka for providing code fore processing eBird data. All data, scripts and full results are available here: https://osf.io/my8pc/

