## Appendix S1 for "Integer linear programming outperforms simulated annealing for solving conservation planning problems"

### Table S1

**Table S1**: List of species that were used as features in our analysis.

| Species Code | Common Name | Scientific Name |
|---|---|---|
| amegfi | American Goldfinch | Spinus tristis |
| amekes | American Kestrel | Falco sparverius |
| amerob | American Robin | Turdus migratorius |
| annhum | Anna's Hummingbird | Calypte anna |
| baleag | Bald Eagle | Haliaeetus leucocephalus |
| barswa | Barn Swallow | Hirundo rustica |
| brdowl | Barred Owl | Strix varia |
| belkin1 | Belted Kingfisher | Megaceryle alcyon |
| bewwre | Bewick's Wren | Thryomanes bewickii |
| bnhcow | Brown-headed Cowbird | Molothrus ater |
| bkhgro | Black-headed Grosbeak | Pheucticus melanocephalus |
| brebla | Brewer's Blackbird | Euphagus cyanocephalus |
| brncre | Brown Creeper | Certhia americana |
| batpig1 | Band-tailed Pigeon | Patagioenas fasciata |
| bushti | Bushtit | Psaltriparus minimus |
| cangoo | Canada Goose | Branta canadensis |
| chbchi | Chestnut-backed Chickadee | Poecile rufescens |
| cedwax | Cedar Waxwing | Bombycilla cedrorum |
| chispa | Chipping Sparrow | Spizella passerina |
| coohaw | Cooper's Hawk | Accipiter cooperii |
| comrav | Common Raven | Corvus corax |
| amecro | American Crow | Corvus brachyrhynchos |
| dowwoo | Downy Woodpecker | Dryobates pubescens |
| eucdov | Eurasian Collared-Dove | Streptopelia decaocto |
| eursta | European Starling | Sturnus vulgaris |
| evegro | Evening Grosbeak | Coccothraustes vespertinus |
| norfli | Northern Flicker | Colaptes auratus |
| foxspa | Fox Sparrow | Passerella iliaca |
| gockin | Golden-crowned Kinglet | Regulus satrapa |
| haiwoo | Hairy Woodpecker | Dryobates villosus |
| houfin | House Finch | Haemorhous mexicanus |
| houspa | House Sparrow | Passer domesticus |
| houwre | House Wren | Troglodytes aedon |
| hutvir | Hutton's Vireo | Vireo huttoni |
| macwar | MacGillivray's Warbler | Geothlypis tolmiei |
| moudov | Mourning Dove | Zenaida macroura |
| norhar1 | Hen Harrier | Circus cyaneus |
| orcwar | Orange-crowned Warbler | Oreothlypis celata |
| olsfly | Olive-sided Flycatcher | Contopus cooperi |
| osprey | Osprey | Pandion haliaetus |
| pacwre1 | Pacific Wren | Troglodytes pacificus |
| pinsis | Pine Siskin | Spinus pinus |
| pilwoo | Pileated Woodpecker | Dryocopus pileatus |
| pasfly | Pacific-slope Flycatcher | Empidonax difficilis |
| purfin | Purple Finch | Haemorhous purpureus |
| purmar | Purple Martin | Progne subis |
| rebnut | Red-breasted Nuthatch | Sitta canadensis |
| rebsap | Red-breasted Sapsucker | Sphyrapicus ruber |
| redcro | Red Crossbill | Loxia curvirostra |

| Species Code | Common Name | Scientific Name |
|---|---|---|
| rocpig | Rock Pigeon | Columba livia |
| rethaw | Red-tailed Hawk | Buteo jamaicensis |
| rufhum | Rufous Hummingbird | Selasphorus rufus |
| rewbla | Red-winged Blackbird | Agelaius phoeniceus |
| savspa | Savannah Sparrow | Passerculus sandwichensis |
| sora | Sora | Porzana carolina |
| sonspa | Song Sparrow | Melospiza melodia |
| spotow | Spotted Towhee | Pipilo maculatus |
| stejay | Steller's Jay | Cyanocitta stelleri |
| swathr | Swainson's Thrush | Catharus ustulatus |
| towwar | Townsend's Warbler | Setophaga townsendi |
| treswa | Tree Swallow | Tachycineta bicolor |
| daejun | Dark-eyed Junco | Junco hyemalis |
| yerwar | Yellow-rumped Warbler | Setophaga coronata |
| varthr | Varied Thrush | Ixoreus naevius |
| vigswa | Violet-green Swallow | Tachycineta thalassina |
| warvir | Warbling Vireo | Vireo gilvus |
| whcspa | White-crowned Sparrow | Zonotrichia leucophrys |
| westan | Western Tanager | Piranga ludoviciana |
| wilsni1 | Wilson's Snipe | Gallinago delicata |
| wlswar | Wilson's Warbler | Cardellina pusilla |
| wooduc | Wood Duck | Aix sponsa |
| yelwar | Yellow Warbler | Setophaga petechia |

**Figure S1**

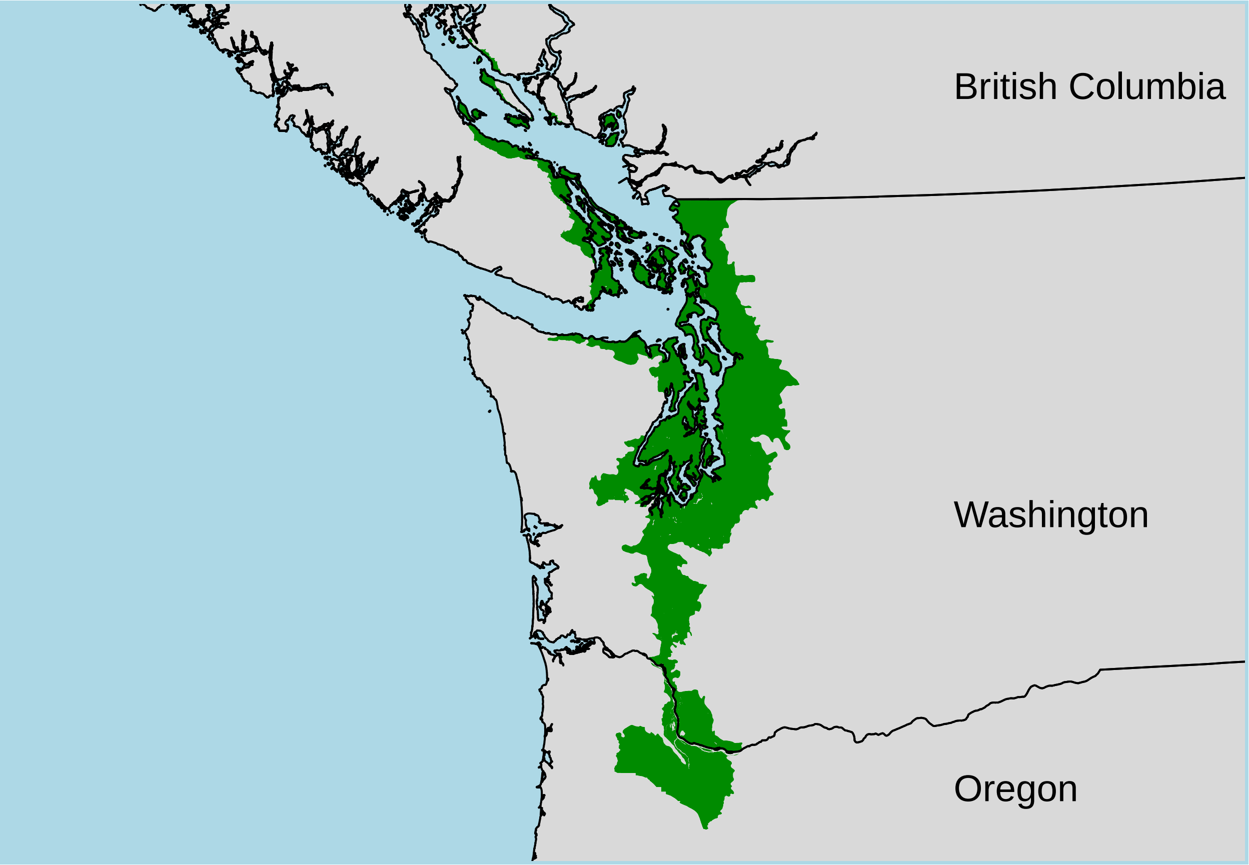


**Figure S1**: Study area.

**Figure S2**

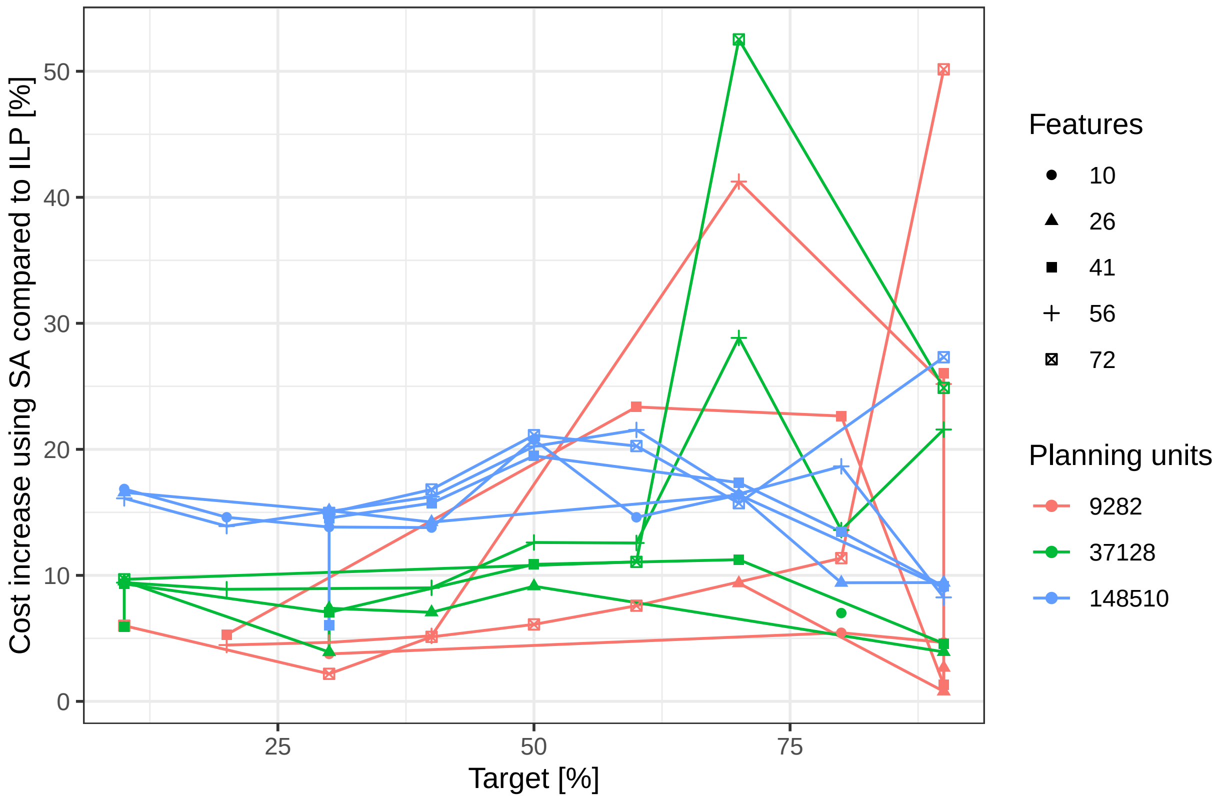


**Figure S2**: Percent cost increase of SA solutions compared to ILP solutions, across targets, number of features and number of planning units. Simulated annealing (i.e. Marxan) parameters used are: number of iterations > 100,000; species penalty factor 5 or 25. Not all Marxan scenarios generated yielded feasible solutions (where all targets were met), which is why e.g. there is only one observation for 37,128 planning units and 10 features.

### Figure S3

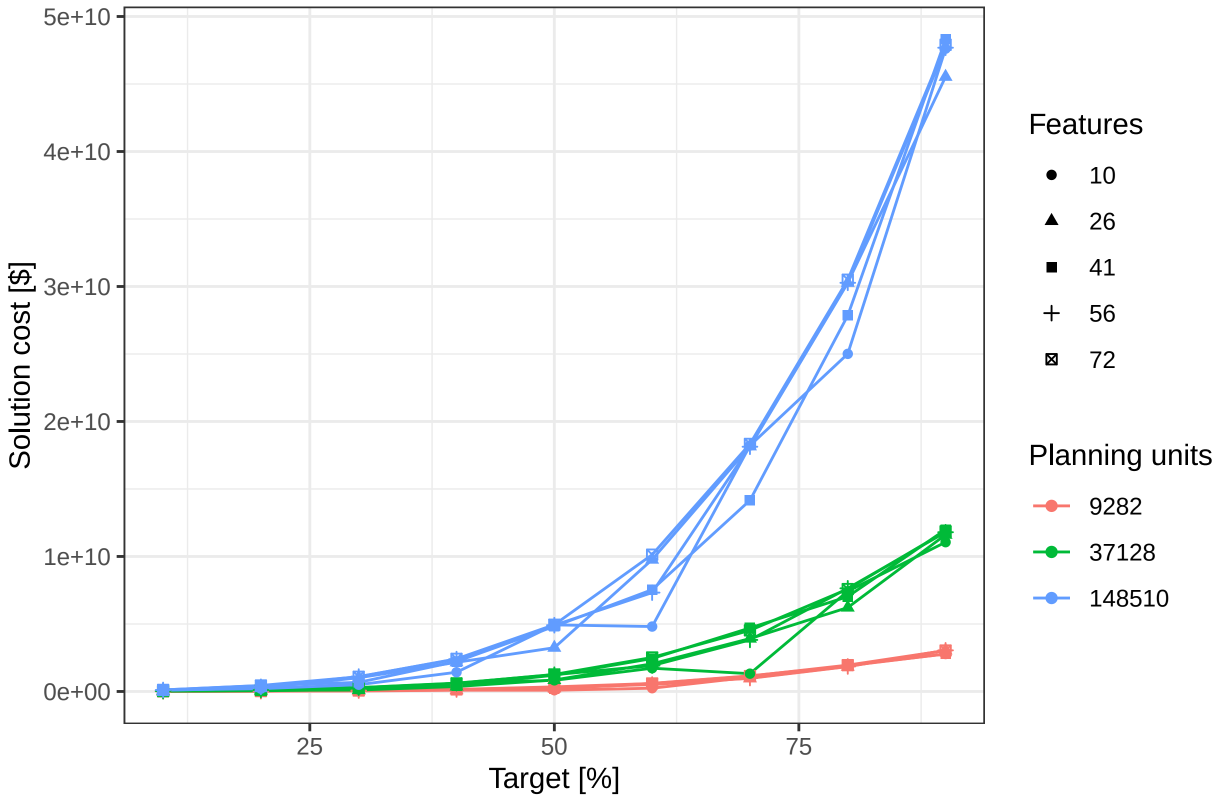


**Figure S3**: Cost profile for Gurobi solver across targets, number of features and number of planning units.

### Figure S4

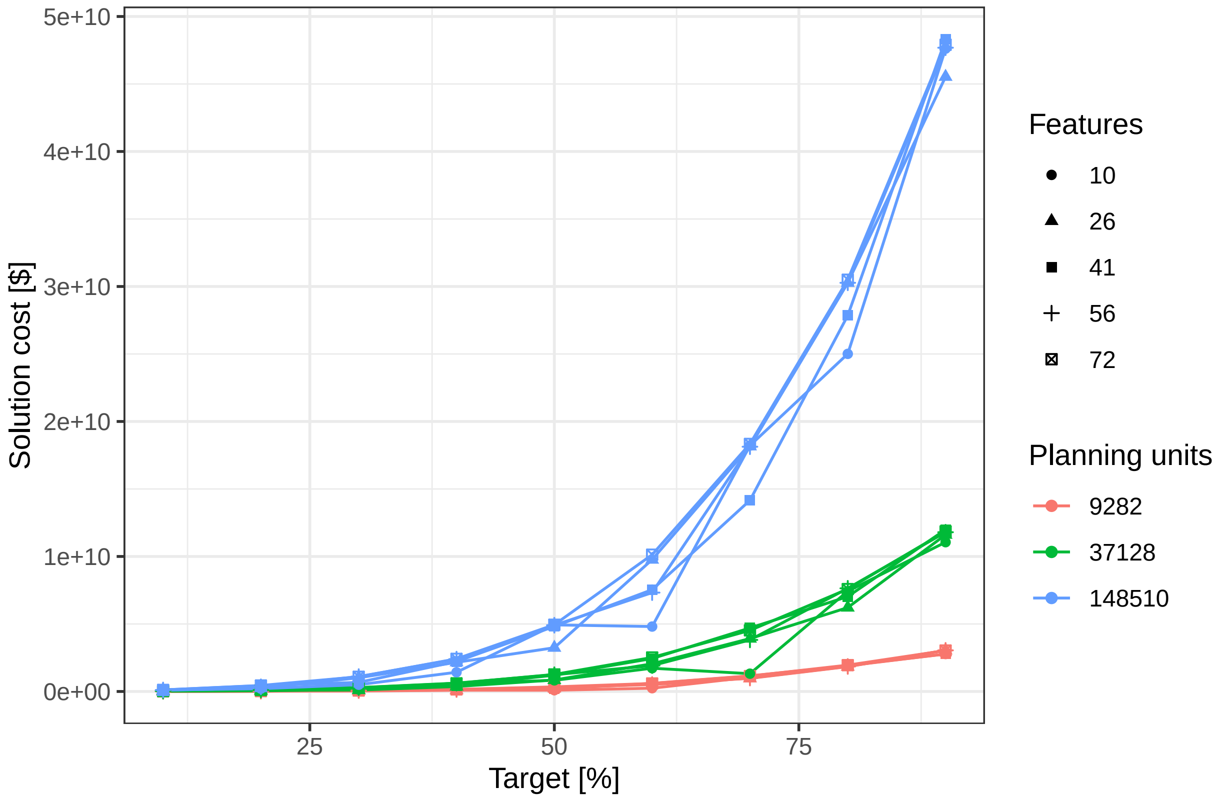


**Figure S4**: Cost profile for SYMPHONY solver across targets, number of features and number of planning units.

**Figure S5**

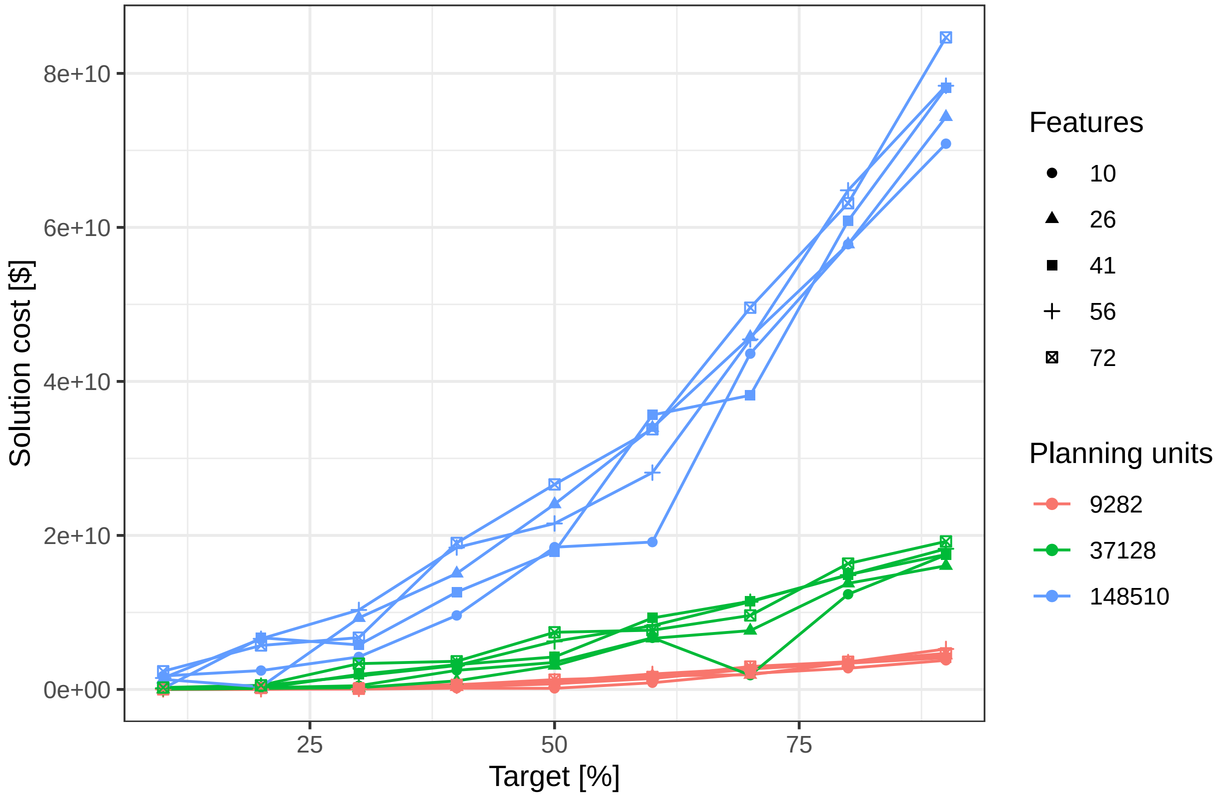


**Figure S5**: Cost profile for Marxan using Simulated Annealing across targets, number of features and number of planning units.

### Figure S6

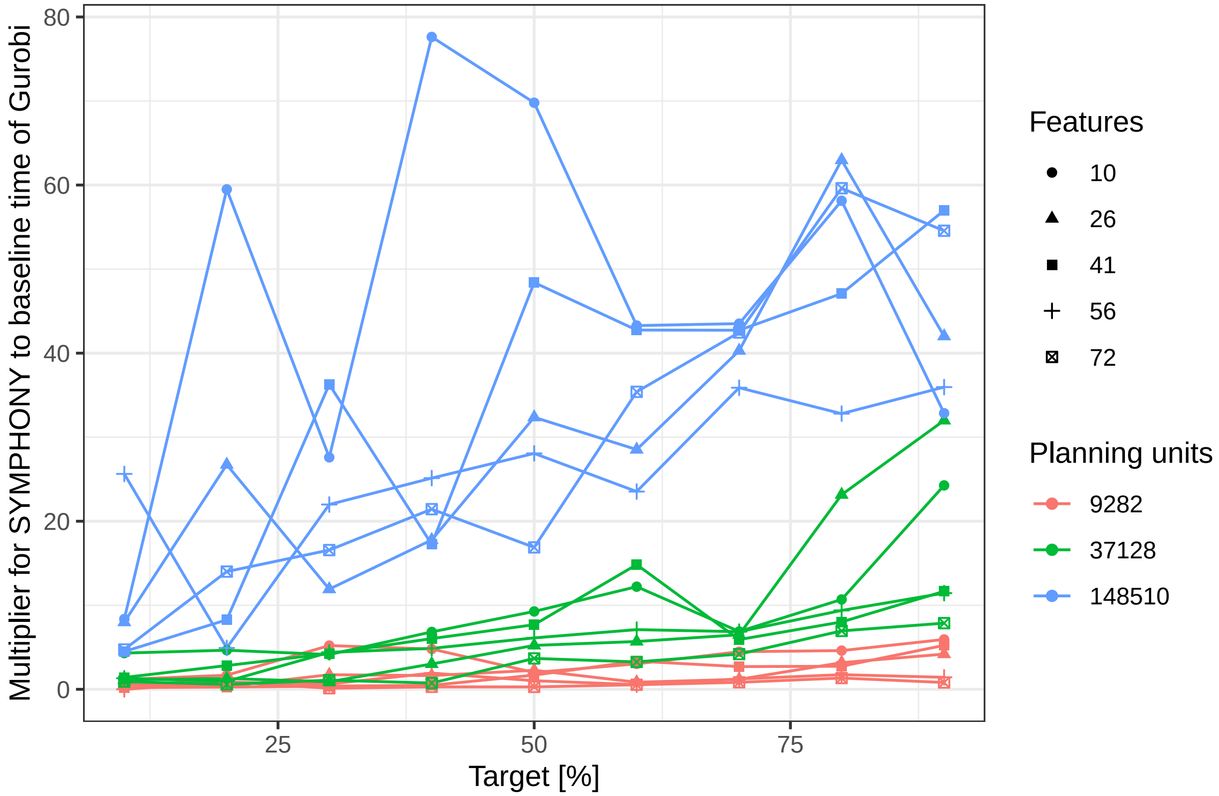


**Figure S6**: Time to solution comparisons between SYMPHONY and Gurobi across targets, number of features and number of planning units.

**Figure S7**

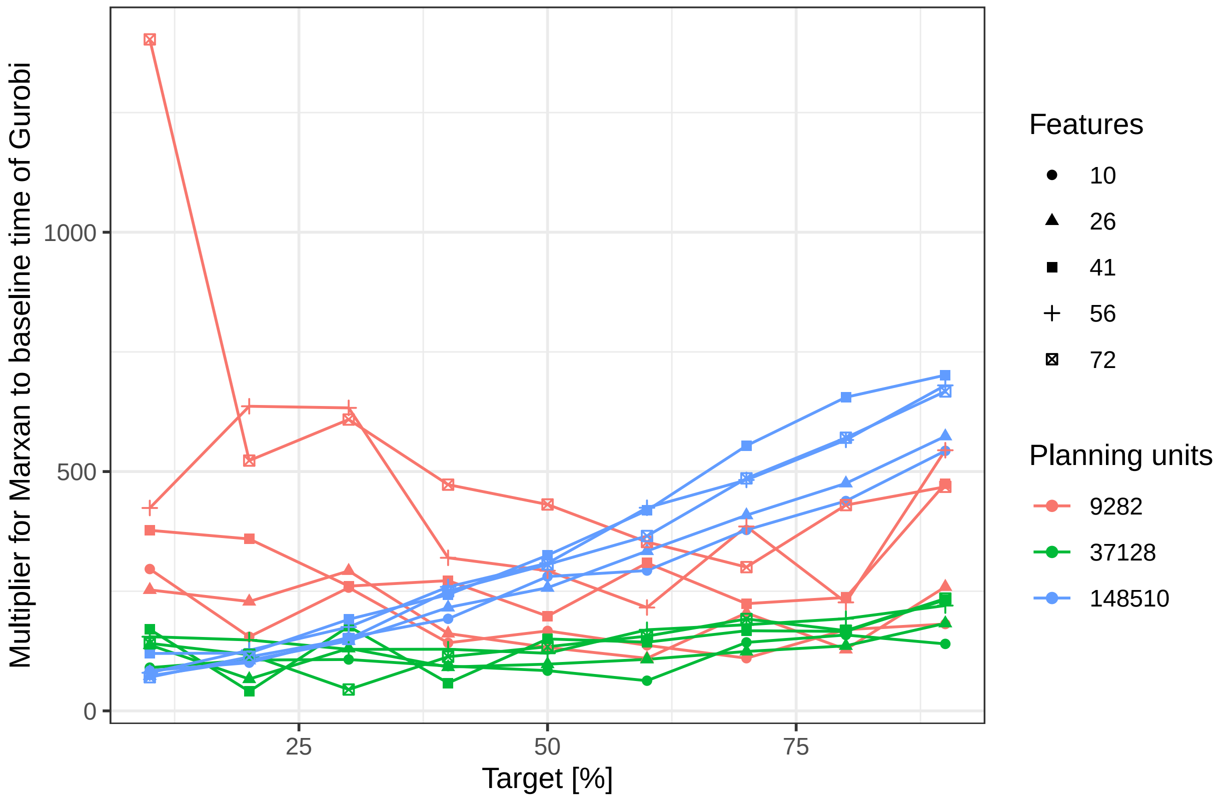


**Figure S7**: Time to solution comparisons between Marxan using Simulated Annealing and Gurobi across targets, number of features and number of planning units.

### Figure S8

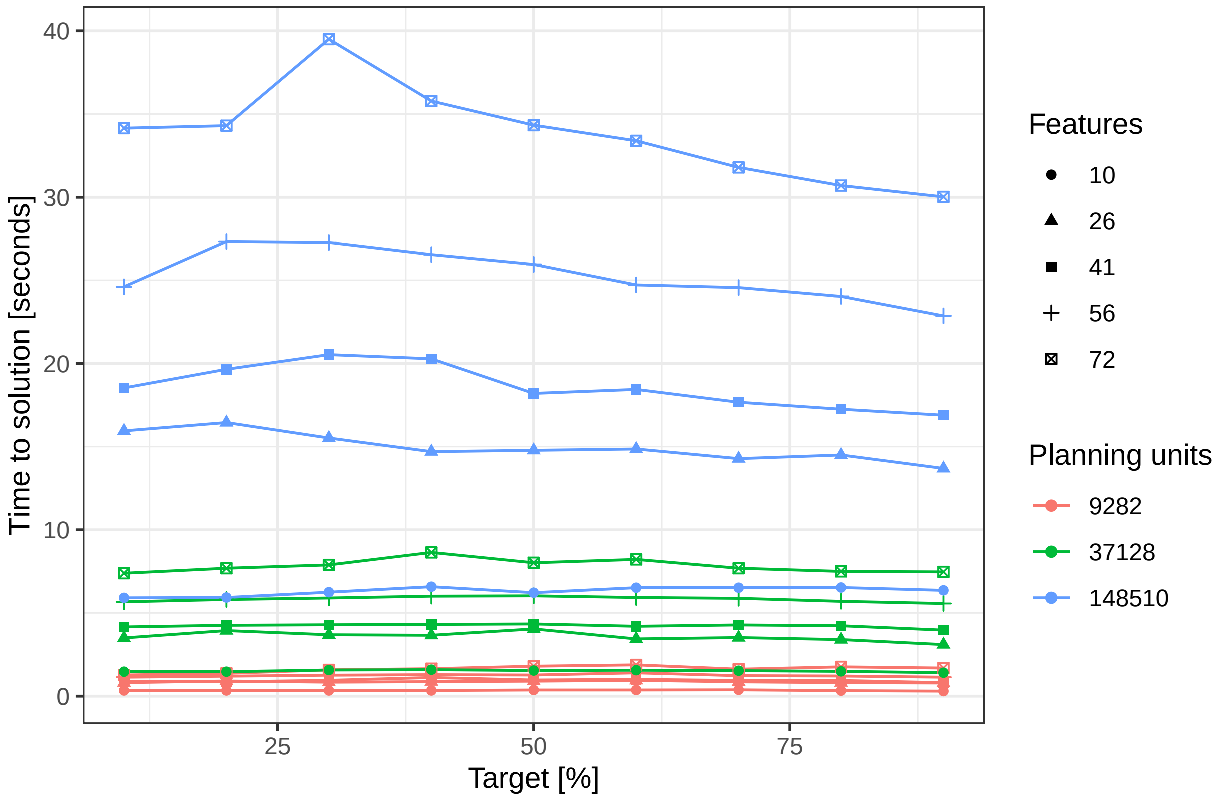


**Figure S8**: Time to solution profile for Gurobi solver across targets, number of features and number of planning units.

**Figure S9**

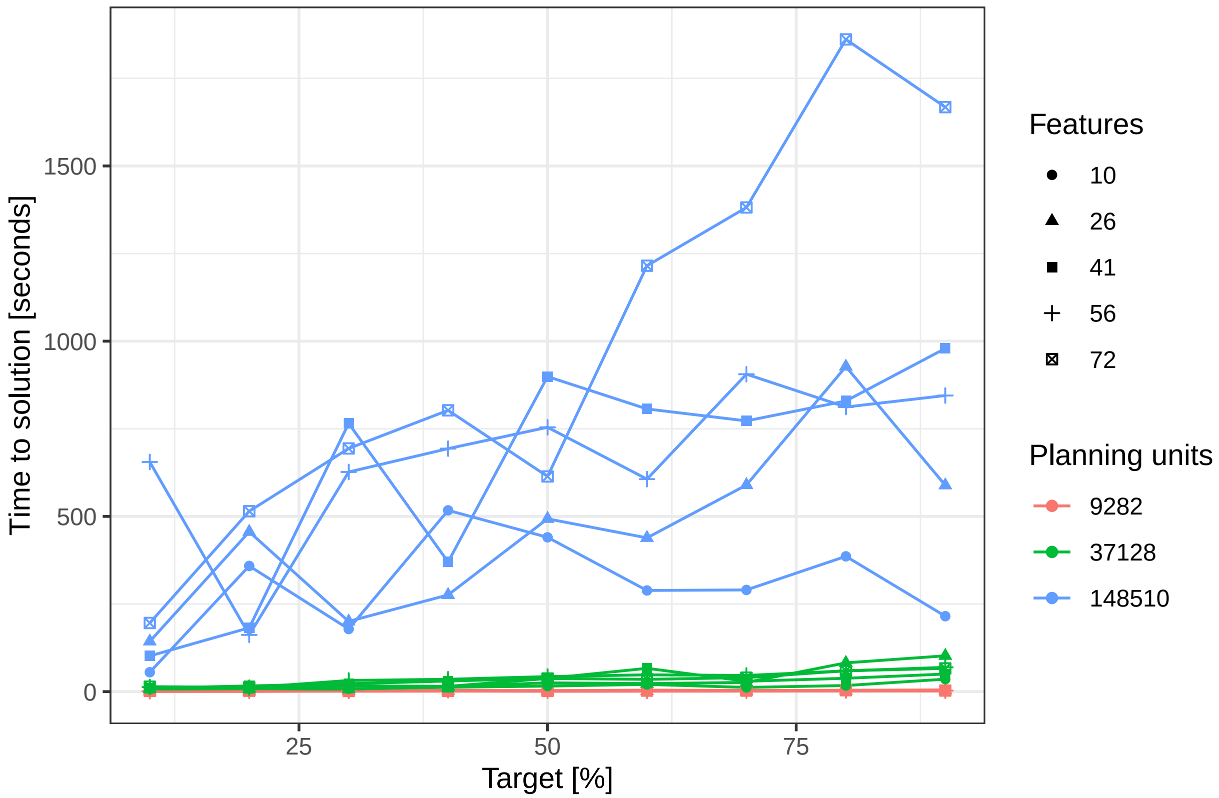


**Figure S9**: Time to solution profile for SYMPHONY solver across targets, number of features and number of planning units.

**Figure S10**

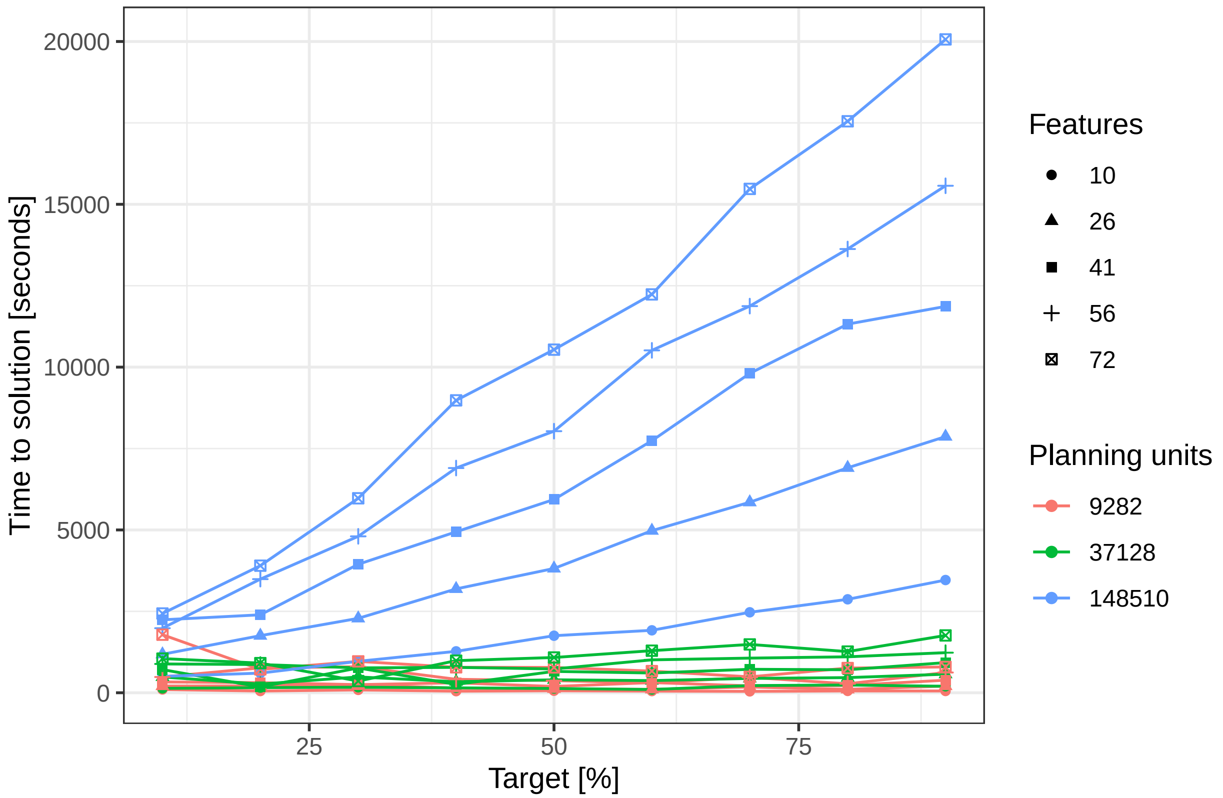


**Figure S10**: Time to solution profile for Marxan using Simulated Annealing across targets, number of features and number of planning units.

**Figure S11**

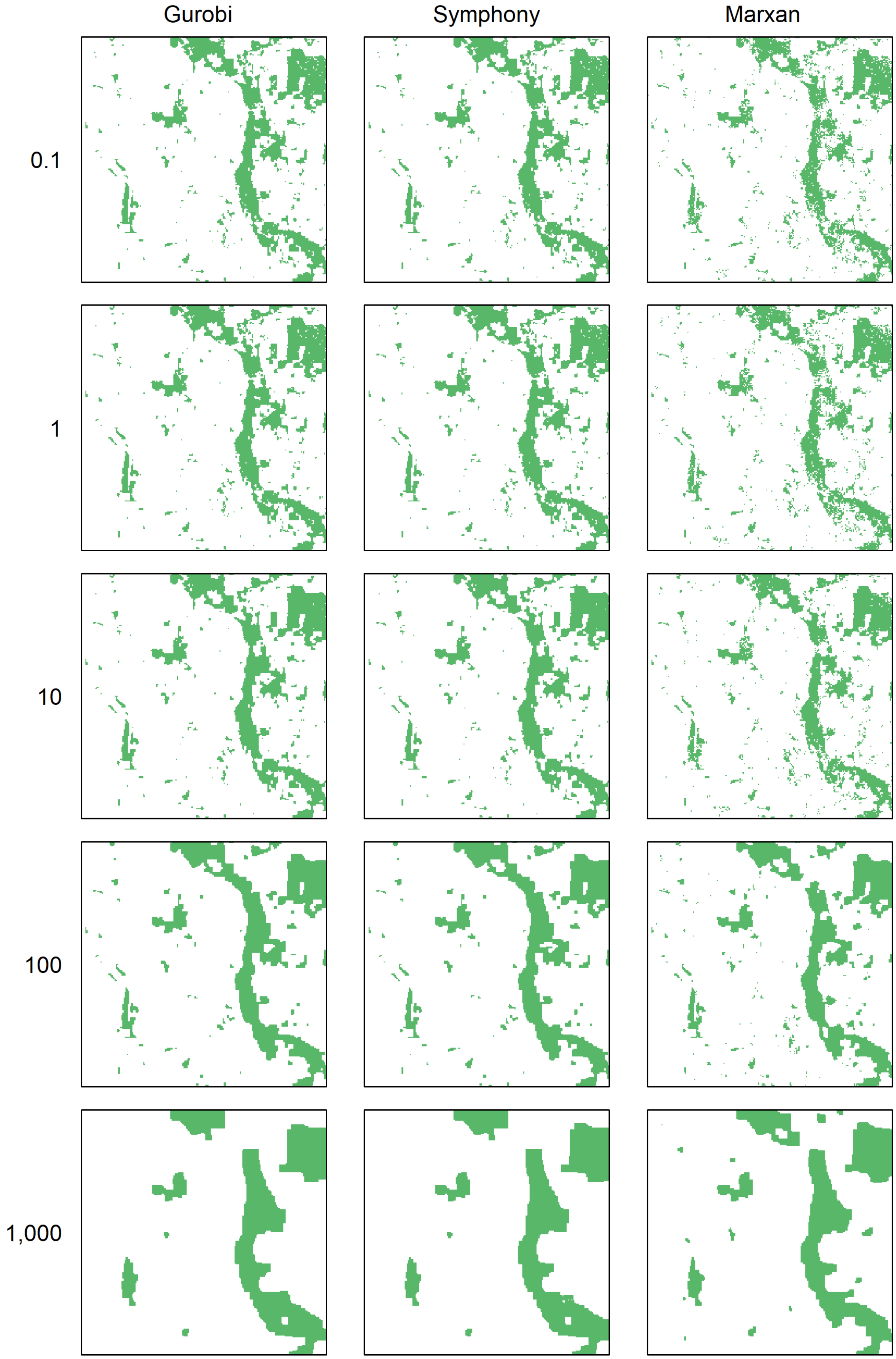


**Figure S11**: Compactness of solutions. Shown are the solutions for a 10% target. The numbers represent BLM values.
